# Mitoquinone mesylate targets SARS-CoV-2 infection in preclinical models

**DOI:** 10.1101/2022.02.22.481100

**Authors:** Anton Petcherski, Madhav Sharma, Sandro Satta, Maria Daskou, Hariclea Vasilopoulos, Cristelle Hugo, Eleni Ritou, Barbara Jane Dillon, Eileen Fung, Gustavo Garcia, Claudio Scafoglio, Arunima Purkayastha, Brigitte N Gomperts, Gregory A Fishbein, Vaithilingaraja Arumugaswami, Marc Liesa, Orian S Shirihai, Theodoros Kelesidis

**Author notes:** Equal authorship (Anton Petcherski, Madhav Sharma, Sandro Satta). These authors contributed equally to this work (Anton Petcherski, Madhav Sharma).

## Abstract

To date, there is no effective oral antiviral against SARS-CoV-2 that is also anti-inflammatory. Herein, we show that the mitochondrial antioxidant mitoquinone/mitoquinol mesylate (Mito-MES), a dietary supplement, has potent antiviral activity against SARS-CoV-2 and its variants of concern *in vitro* and *in vivo*. Mito-MES had nanomolar *in vitro* antiviral potency against the Beta and Delta SARS-CoV-2 variants as well as the murine hepatitis virus (MHV-A59). Mito-MES given in SARS-CoV-2 infected K18-hACE2 mice through oral gavage reduced viral titer by nearly 4 log units relative to the vehicle group. We found *in vitro* that the antiviral effect of Mito-MES is attributable to its hydrophobic dTPP+ moiety and its combined effects scavenging reactive oxygen species (ROS), activating Nrf2 and increasing the host defense proteins TOM70 and MX1. Mito-MES was efficacious reducing increase in cleaved caspase-3 and inflammation induced by SARS-CoV2 infection both in lung epithelial cells and a transgenic mouse model of COVID-19. Mito-MES reduced production of IL-6 by SARS-CoV-2 infected epithelial cells through its antioxidant properties (Nrf2 agonist, coenzyme Q10 moiety) and the dTPP moiety. Given established safety of Mito-MES in humans, our results suggest that Mito-MES may represent a rapidly applicable therapeutic strategy that can be added in the therapeutic arsenal against COVID-19. Its potential *long-term* use by humans as diet supplement could help control the SARS-CoV-2 pandemic, especially in the setting of rapidly emerging SARS-CoV-2 variants that may compromise vaccine efficacy.

**One-Sentence Summary:** Mitoquinone/mitoquinol mesylate has potent antiviral and anti-inflammatory activity in preclinical models of SARS-CoV-2 infection.

## Main Text

The SARS-CoV-2 pandemic emphasizes the urgent need to determine cellular pathways that can be targeted by novel oral antivirals that ideally would target rapidly evolving SARS-CoV-2 variants *and* associated inflammation that drives morbidity in COVID-19(*1*). Current oral antivirals against SARS-CoV-2 such as nirmatrelvir/ritonavir (paxlovid) and molnupiravir do not have anti-inflammatory effects. Molnupiravir may be mutagenic in cells(*2*). COVID-19 rebound has been described after use of paxlovid which may cause life-threatening interactions with widely used medications. Thus, there is a continued need for the development of oral safe antivirals for COVID-19.

Increased reactive oxygen species (ROS) contribute to pathogenesis of respiratory syncytial virus (RSV)(*3*) and coronavirus infection(*4*). Mitochondria are a major source of ROS (mito-ROS). Mitoquinone and/or mitoquinol mesylate (Mito-MES) is the only mitochondrial antioxidant approved for human use which has safely been used in clinical trials for ROS-related diseases (*5*, *6*). Mito-MES is coenzyme Q_10_ (CoQ_10_) conjugated to a lipophilic cation (TPP)(*5*). CoQ_10_ is the endogenous mitochondrial coenzyme involved in electron transfer and protection from lipid peroxidation(*5*). Given that Mito-MES is antiviral against RSV(*3*), we investigated whether Mito-MES is antiviral against SARS-CoV-2.

We first determined the effects of Mito-MES on SARS-CoV-2 replication in human lung epithelial Calu-3 (Calu3) cells. In non-cytotoxic concentrations (based on XTT assay), the half maximal inhibitory concentration (IC50) of Mito-MES against SARS-CoV-2 measured by TCID50 was 30-fold lower than remdesivir (Fig. 1A and B). qPCR (Fig. S1A), immunofluorescence (IF) (Fig. 1C) and flow cytometry (FC) (Fig. S1B) also showed that Mito-MES had antiviral activity in Calu3 and in human airway epithelial (HAE) cells in air-liquid interface (ALI) cultures (Fig. 1D). The anti-SARS-CoV-2 activity of Mito-MES was higher in human than Vero-E6 cells as shown by qPCR (Fig. S1C), IF (Fig. S1D), FC (Fig. S1E) and TCID50 assay (Fig. S1F). Notably, TCID50 assays showed that unlike remdesivir (Fig. S1G) the IC50 of Mito-MES was ~100-fold lower in Calu3 than Vero-E6 cells (Fig. 1A and Fig. S1F). Similarly, TCID50 assays showed that the anti-SARS-CoV-2 activity of Mito-MES was higher in human cells than in mouse lung cells from K18-hACE2 mice (Fig. S1H) and higher than remdesivir (Fig. S1I). We found that the antiviral activity of Mito-MES was non-linear between different doses (Fig. S1A and C and D) and was more potent in lower rather than higher concentrations in Calu3 cells (Fig. S1A). Overall, our data suggest that unlike other antivirals that target the viral replication machinery and have dose-dependent antiviral effects, Mito-MES acts through non-linear host antiviral mechanisms that are species-dependent.

**Fig.1.**
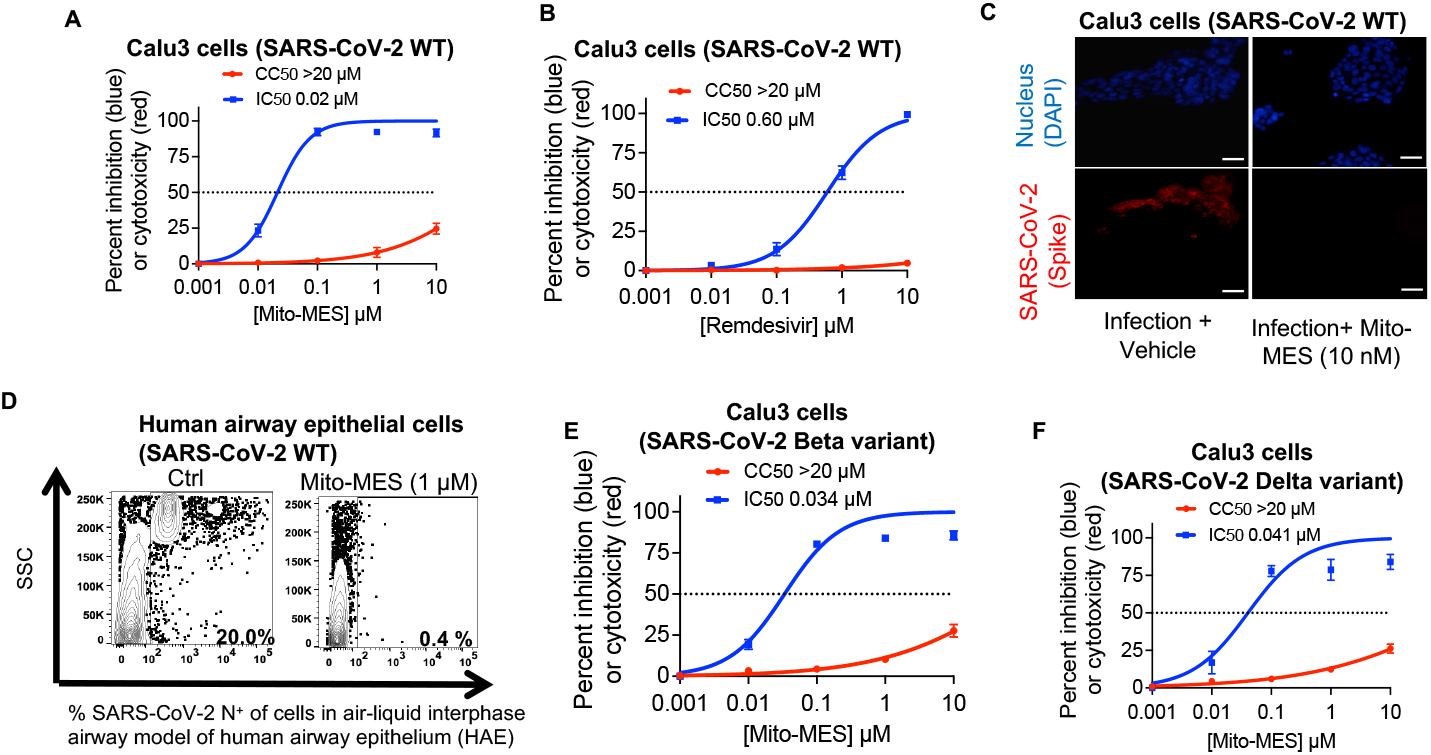
Mitoquinone mesylate (Mito-MES) has strong anti-SARS-CoV-2 activity in multiple human airway cells and has additive antiviral effects with paxlovid. Calu3 cells, human airway epithelial cells cultured in air-liquid interface and high ACE2 (hACE2) A549 cells were infected with wild type (WT) SARS-CoV-2 or B.1.351 (Beta), B.1.617.2 (Delta), B.1.1.529 (Omicron) variants at an MOI of 0.1 (48 hrs) and were treated with drugs for at least 1 hour before infection and throughout the experiment. Viral replication at 48 hours post infection (hpi) by TCID50-assay or flow cytometry. Cell cytotoxicity was assessed in uninfected cells by the XTT (48 h). IC50, 50% cytotoxic concentration (CC50) values for drugs [Mito-MES, remdesivir (RDV)] are indicated. **(C)** Immunofluorescent analysis (48 hpi) of infected Calu3 cells. Scale bar=50μm. In all panels, data are means ± SEM of at least three independent experiments with at least 3 replicates in each experiment.

We hypothesized that Mito-MES has host antiviral activity against SARS-CoV-2 variants of concerns (VOCs) and other coronaviruses. TCID50 assays showed that Mito-MES inhibited *in vitro* SARS-CoV-2 B.1.351 (Beta) (Fig. 1E) and B.1.617.2 (Delta) variants (Fig. 1F) in Calu3 cells. Finally, Mito-MES also inhibited murine hepatitis virus (MHV-A59) in mouse 17Cl-1 fibroblasts (Fig. S1J).

To elucidate the mechanism of antiviral action of Mito-MES, we compared the antiviral effects of Mito-MES administered before or concurrently with SARS-CoV-2. We found that 1000 nM (concentration that was consistently antiviral among all different experimental systems) Mito-MES inhibited SARS-CoV-2 nucleocapsid protein (NP) expression in Calu3 cells when added before infection (effect on viral entry) but also when added 4 hours post infection (hpi), which allows time for viral entry, in both Calu3 (Fig. 2A) and hACE2-HEK293T cells (Fig. 2B) infected with Beta variant. Thus, Mito-MES inhibits both viral entry and cytoplasmic replication.

**Fig.2.**
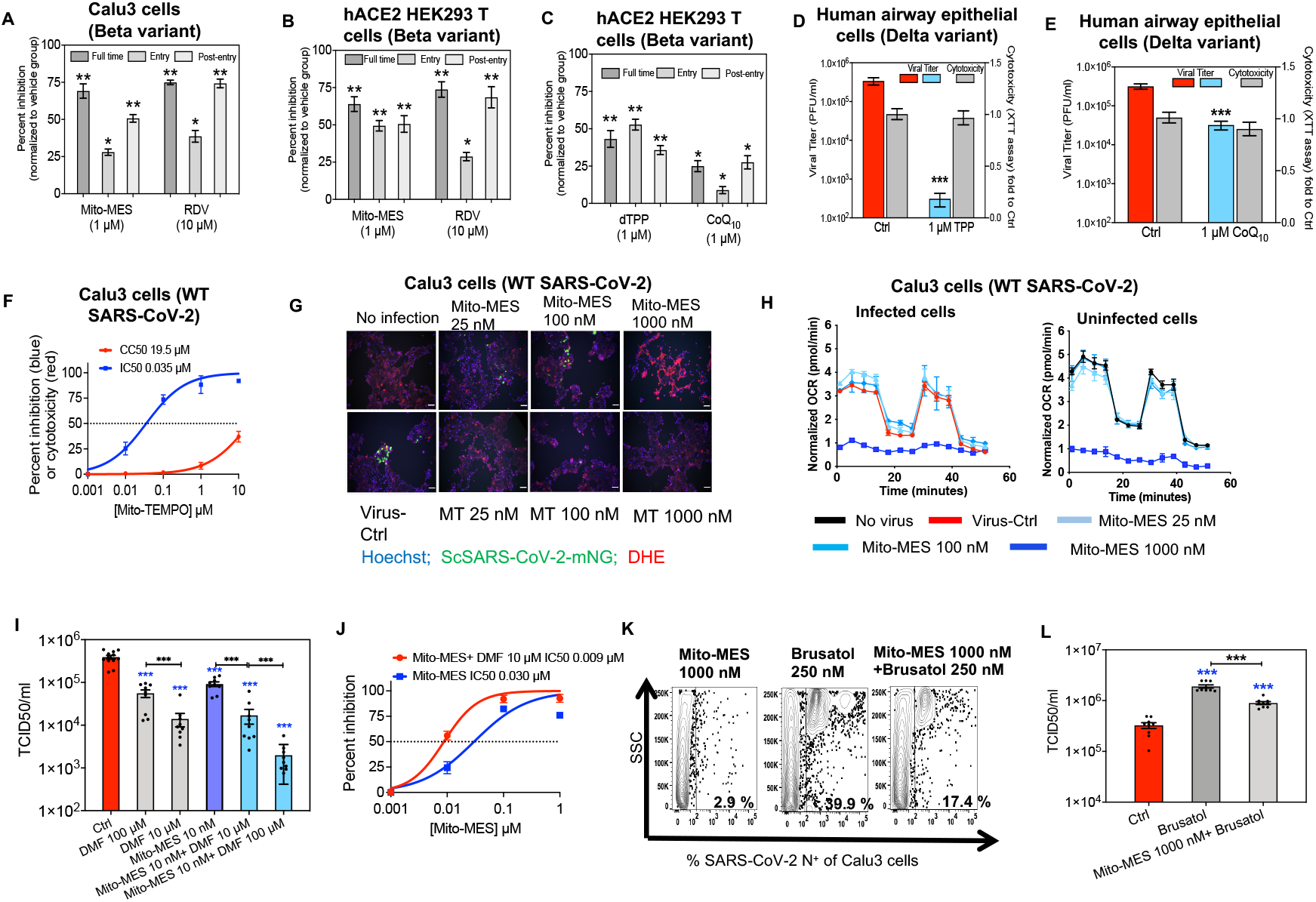
Mitoquinone mesylate (Mito-MES) has anti-SARS-CoV-2 activity through its lipophilic and antioxidant components. (**A** to **C**) Time-of-addition experiment of remdesivir (RDV), Mito-MES, dTPP and coenzyme Q10 (CoQ_10_) in Calu3 and hACE2 HEK293T cells [at entry, post-viral entry and throughout 24 hpi (full time)]. Levels of intracellular SARS-CoV-2 Nucleocapsid protein (NP) were determined by ELISA. (**D** to **L**) Human airway epithelial and Calu3 cells were infected and treated as shown. Viral replication by TCID50-assay and flow cytometry (**K**) and XTT assay at 48 hpi. IC50 and CC50 values are indicated. **(G**) Oxidative stress by dihydroethidum (DHE) fluorescence and replication of fluorescent icSARS-CoV-2-mNG (MOI 0.3) at 48 hpi by immunofluorescence. **(H**) Seahorse Analyzer determined oxygen consumption rate (OCR) in (un)infected Calu3 cells. For all experiments except (**G**) cells were infected with SARS-CoV-2 and treated as in Fig. 1. In all panels, data are representative or mean ± SEM of at least two experiments in 3 replicates. Statistical comparison was done between the Ctrl and each group (n=5-10/group) by two-tailed Mann–Whitney (\**p*□<□0.05, \*\**p*□<□0.01, \*\*\**p*□<□0.001). For “Entry” treatment, the drugs were added to the cells for 2□h before and after infection and the supernatant was then replaced with culture medium.

The lipophilic TPP moiety of Mito-MES facilitates entry of CoQ_10_ through phospholipid bilayers so that it can scavenge ROS more efficiently(*7*). Given that membrane lipid layers and mito-ROS are important for SARS-CoV-2 pathogenesis, we hypothesized that the lipophilic TPP and CoQ_10_ moieties of Mito-MES have anti-SARS-CoV-2 activity. We confirmed by TCID50 assays in hACE2-HEK293T cells and HAE cultures that non-cytotoxic 1000 nM TPP and CoQ_10_ had potent antiviral activity against the Beta and Delta variants (Fig. 2C to E). These data showed that the antiviral activity of Mito-MES in epithelial cells is partially mediated through its lipophilic components.

SARS-CoV2 infection increases mito-ROS(*4*) and the antiviral activity of Mito-MES against RSV is mainly achieved by decreasing mito-ROS (*3*). We hypothesized that the antioxidant activity of Mito-MES contributes to its anti-SARS-CoV-2 activity. Mito-MES reduced SARS-CoV-2-induced increase in fluorescence of the mitochondrial superoxide reporter MitoSOX Red assessed by FC (Fig. S2A to D). Both Mito-TEMPO, an independent mitochondrial antioxidant that also has TPP(*7*), and Mito-MES had potent and similar anti-SARS-CoV-2 activity in Calu3 cells (Fig. 2F and G) and Vero-E6 cells (Fig. S2E). Notably, consistent with prior evidence that high concentrations of Mito-MES can have *in vitro* prooxidant effect that does not appear *in vivo*(*5*), treatment with 1000 nM Mito-MES increased ROS levels in Calu3 cells, while preserving potent antiviral activity (Fig. 2G). Given that mito-ROS regulate mitochondrial function, we interrogated the impact of SARS-CoV-2 on cellular bioenergetics and found that SARS-CoV-2 reduced oxygen consumption rate (OCR) and increased extracellular acidification rate (ECAR) in Calu3 cells (Fig. S2F and G). Consistent with prior data that excessive ROS scavenging decreases mitochondrial function(*8*), antiviral doses of Mito-MES 50-1000 nM reduced OCR (Fig. 2H) and ECAR (Fig. S2H) *in vitro* regardless of SARS-CoV-2 infection. Consequently, our data supported that the anti-SARS-CoV-2 activity of Mito-MES may be mediated through *non-mitochondrial* host pathways.

Mito-MES induces the antiviral antioxidant Nrf2 pathway in epithelial cells(*9*). We determined whether Mito-MES has Nrf2-mediated antiviral activity. Pretreatment of Calu3 with the Nrf2 agonist Dimethyl fumarate (DMF) for 24 hours increased in a dose dependent manner the anti–SARS-CoV-2 activity of Mito-MES by TCID50 (Fig. 2I and J). Pretreatment of Calu3 for 24 hours with the Nrf2 inhibitor brusatol (0.25 μM)(*10*) significantly increased SARS-CoV2 replication as assessed by FC and TCID50 and inhibited the antiviral effect of Mito-MES (Fig. 2K and L). 100 μM DMF, but not 0.25 μM brusatol and 10 μM DMF, were cytotoxic in uninfected cultures (Fig. S3A). Flow cytometry showed that Mito-MES did not impact levels of Nrf2 (Fig. S3B) but reduced levels of the endogenous Nrf2 inhibitor, Keap1 (Fig. S3C) and increased levels of heme oxygenase-1 (HO-1) (Fig. S3D), a key antiviral protein of the Nrf2 pathway(*11*), in (un)infected Calu3 cells. Overall, our data suggested that Mito-MES has Nrf2-mediated anti-SARS-CoV-2 activity and are in agreement with Olagnier et al who showed that a cytotoxic (*10*) concentration of DMF (200 μM) had anti-SARS-CoV2 activity (*12*).

Given that Mito-MES was less antiviral in interferon deficient Vero-E6 cells compared to human cells (Fig. 1), we hypothesized that interferon pathways that interact with mitochondrial proteins such as MAVS and TOM70(*13*, *14*), mediate its antiviral activity. Using FC and IF in (un)infected Calu3 cells, we found that Mito-MES had no impact on MAVS (Fig. S3E) but increased TOM70 (Fig. S3F and G) and MX1 (Fig. S3H), a key antiviral effector in COVID-19 patients(*15*) that interacts with mitochondria(*16*). ELISA showed that at 48 hpi, Mito-MES did not impact secretion of IFN-β and IFN-*λ* by SARS-CoV-2 infected Calu3 cells (Fig. S3I). Overall, our data demonstrate that Mito-MES selectively induces mitochondrial mediators of IFN-I which can explain better antiviral activity in interferon competent cells.

Mito-MES has also anti-inflammatory effects (*5*). Thus, we also assessed the impact of Mito-MES on SARS-CoV-2-associated inflammatory responses that contribute to development of severe COVID-19 such as increased IL-1β and IL-6(*1*). Using ELISA and Luminex immunoassays, we found that Mito-MES attenuated SARS-CoV-2-induced increase in secretion of IL-6 by Calu3 (Fig. 3A) cells and of IL-6 and IL-1β in upper (Fig. 3B) and lower (Fig. 3C) airway ALI cultures. Mito-MES has established pharmacokinetics in mice and in humans and good bioavailability at the respiratory mucosa(*5*). Doses of Mito-MES in mice of 4 mg/kg/day intraperitoneally (i.p) and 20 mg/kg/day through gavage (typically higher dose than i.p due to absorption) achieve good lung tissue penetration and have been translated to oral doses in humans(*5*). Therefore, we tested the *in vivo* anti-inflammatory efficacy of 4 mg/kg intraperitoneal daily dosage of Mito-MES in K18-hACE2 mice infected with the SARS-CoV-2 (Fig. 3D to N). Using Luminex immunoassays we showed that there was a reduction of at least 2 orders of magnitude in IL-1β (Fig. 3E) and IL-6 (Fig. 3F) in the lungs of the SARS-CoV-2 infected Mito-MES-treated relative to the vehicle treated mice as early as 3 days post infection (dpi) that was also confirmed after at least 5 dpi in mice infected with the SARS-CoV-2 Beta variant, when lung inflammation is the highest (*17*)(Fig. 3, H and I). Mito-MES also reduced levels of TNF-α (Fig. 3J). Histopathology analysis also showed a reduction of lung inflammation and tissue damage in Mito-MES treated over vehicle-treated mice at 5-7 dpi (Fig. 3K to M).

**Fig.3.**
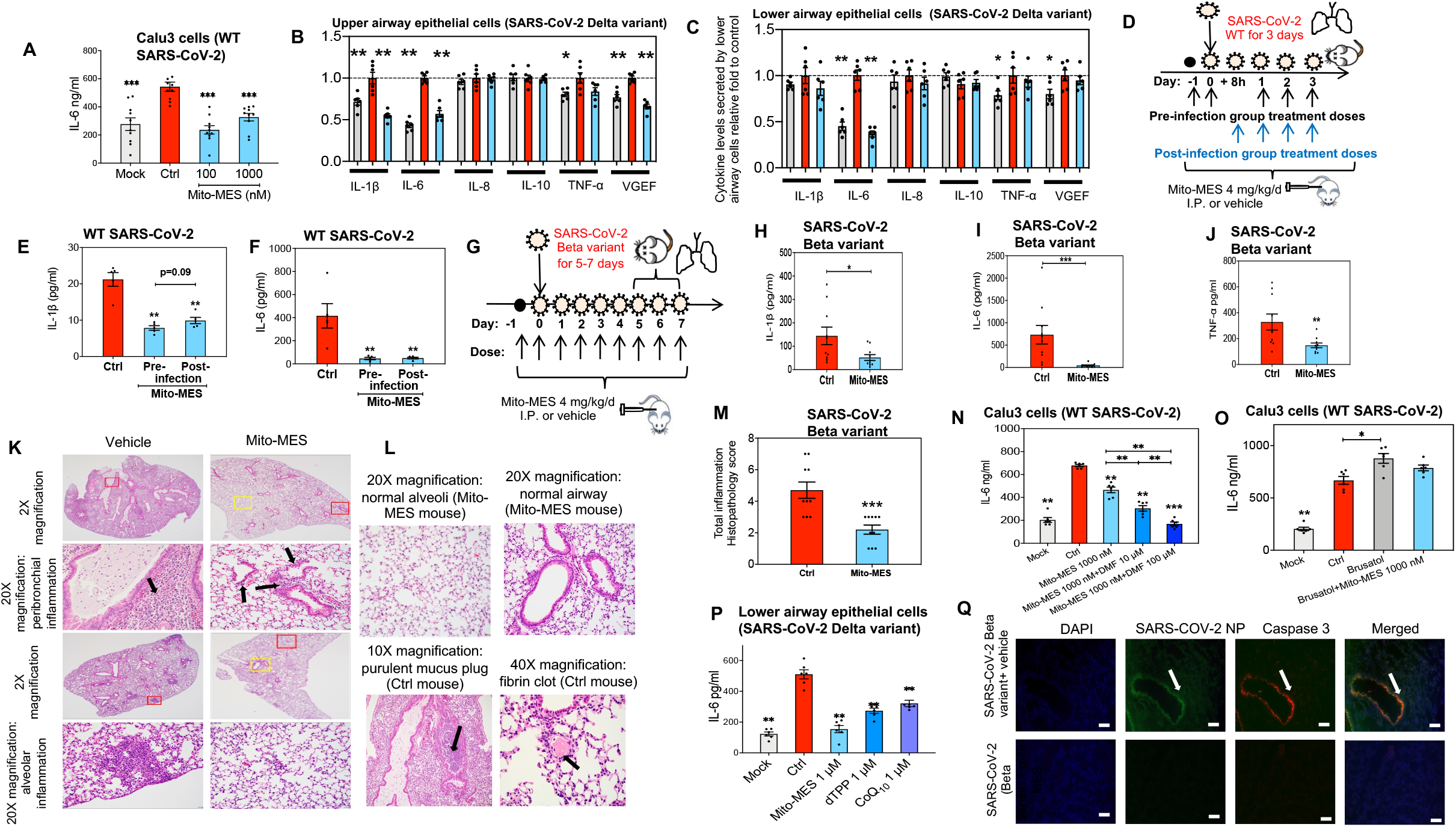
Mitoquinone mesylate (Mito-MES) has anti-inflammatory and anti-apoptotic activity in SARS-CoV-2 infection. (**A** to **R**) Calu3 and airway epithelial cells cultured in air-liquid interface and hACE2 K18 mice were infected (MOI 0.1/cell or 10,000 PFU/mouse) for 72 hours [cells or mice in (**D**)] or at least 5 days [mice in (**G**)(**K**)] with wild type (WT) SARS-CoV-2 or variants (Beta or Delta) and treated as shown. Supernatants from cells or homogenates from harvested lungs were used for measurement of cytokines using Luminex immunoassay or ELISA (**A**). (**L** to **N**) Murine lungs from (**G**) were paraffin-embedded and 5-μm sections were stained for hematoxylin and eosin. Representative images of lung areas where inflammation was assessed are highlighted by red (abnormal area) or yellow (normal area) boxes and arrows. Histopathology score was determined according to Methods (**N**). (**R**) Independent sections were stained with anti-SARS-CoV-2 nucleocapsid (NP) protein (green, white arrow), anti-cleaved caspase 3 (red, white arrow) antibodies and DAPI (blue). Scale bars, 100 μm. Representative or summary (mean□±□SEM) data from experiments in triplicates. Each data-point represents one biological sample. Unless otherwise shown statistical comparison was done between the DMSO (10% with saline in mice) vehicle (Ctrl) and each group (n=5-10/group) by using two-tailed Mann–Whitney ***p<0.001, **p<0.01, *p<0.05.

Nrf2 has a major role in regulation of inflammation and reduces production of IL-1β and IL-6(*11*). We hypothesized that not only the antiviral but also the anti-inflammatory activity of Mito-MES in SARS-CoV-2 infection is Nrf2-mediated. Pretreatment of Calu3 cultures with DMF for 24 hours reduced in an additive and dose dependent manner subsequent release of IL-6 to the cell supernatant from infected cells at 48 hpi as measured by immunoassays (Fig. 3N). Pretreatment of Calu3 with the Nrf2 inhibitor brusatol (*10*) significantly increased subsequent release of IL-6 by infected cells and inhibited the anti-inflammatory effect of Mito-MES (Fig. 3O). Notably, the lipophilic moieties of Mito-MES, dTPP and CoQ_10_ also reduced IL-6 from SARS-CoV-2 infected ALI cultures (Fig. 3P).

Nrf2 pathway and mito-ROS also regulate apoptosis of lung cells(*11*) which is associated with viral replication, lung injury and severe COVID-19(*1*). Thus, we assessed the impact of Mito-MES on SARS-CoV-2 associated cellular apoptosis. Immunostaining showed that Mito-MES reduced SARS-CoV-2-induced increase in cleaved caspase 3 (CC3), a key regulator of apoptosis and cell/tissue damage, in both Vero-E6 (Fig. S4A to E) and Calu3 cells (Fig. S4F to I). Immunofluorescence showed that Mito-MES also inhibited SARS-CoV-2-associated increase in CC3, in K18-hACE2 mice infected with Beta variant (Fig. 3Q). Overall, these data suggest that Mito-MES reduces SARS-CoV-2-induced cell/tissue inflammation and apoptosis through its antioxidant properties (Nrf2 agonist, CoQ_10_ moiety, antioxidant against mito-ROS) and the dTPP.

We then explored the antiviral efficacy of Mito-MES against SARS-CoV-2 in K18-hACE2 mice (*17*). There was a reduction of *at least* 2 log units in SARS-CoV-2 viral titers (Fig. 4A) and SARS-CoV-2 NP protein levels (Fig. 4B) in the lungs of the Mito-MES groups (treated as shown in Fig 3D before and *after* infection) relative to the vehicle control group. The anti-SARS-CoV-2 activity of Mito-MES in K18-hACE2 lung cells was higher than remdesivir (Fig S1H and I) and higher than the previously reported anti-SARS-CoV-2 effect of remdesivir in K18-hACE2 mice (*17*). We then assessed whether oral Mito-MES derived from the formulation given to humans as dietary supplement is antiviral against SARS-CoV-2 *in vivo*. Mito-MES 20 mg/kg/day treatment in SARS-CoV-2 infected (Delta variant) K18-hACE2 mice through oral gavage (treated as shown in Fig.4C) reduced viral titer by nearly 4 log units relative to the vehicle group (Fig. 4D).

**Fig 4.**
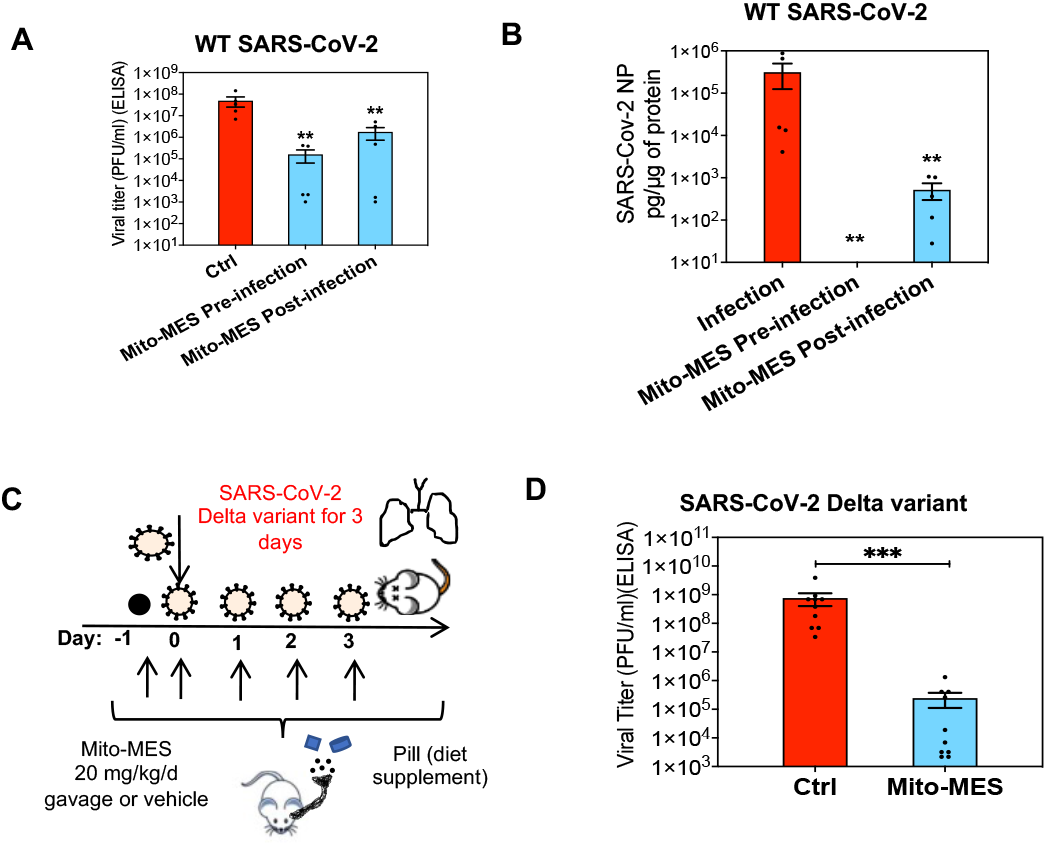
Mitoquinone mesylate (Mito-MES) has anti-SARS-CoV-2 activity in mouse model of SARS-CoV-2 infection. (**A** to **D**) hACE2 K18 mice were infected intranasally with wild type (WT) SARS-CoV-2 (Cohort A; Fig. 3) or Delta variant (**C**) (10,000 PFU/mouse) and treated as shown in Fig. 3D or through gavage as in (**C**). Lung was harvested 72 hours post infection (hpi). ELISA that measures SARS-CoV-2 nucleocapsid protein (NP) was used to determine viral replication in lung protein homogenates (**B**) or in Vero-E6 cells that were used for viral titer in supernatants from murine lung homogenates (**A**). Each data-point represents one biological sample. Unless otherwise shown statistical comparison was done between the DMSO (10% with saline in mice) vehicle (Ctrl) and each group (n=5) by using two-tailed Mann– Whitney, **p<0.01, *p<0.05.

In summary, we demonstrate that Mito-MES has *in vitro* and *in vivo* antiviral, antiapoptotic and anti-inflammatory effects in SARS-CoV-2 infection. Mito-MES may have multiple favorable therapeutic effects in COVID-19 (Fig. S5). We show that the antiviral effect of Mito-MES against SARS-CoV-2 is mediated partially through the Nrf2 pathway, the hydrophobic dTPP cation and interferon responses. Mito-MES had nanomolar antiviral potency against the Beta, Delta SARS-CoV-2 variants as well as MHV. Thus, Mito-MES is expected to have antiviral activity against rapidly emerging SARS-CoV-2 variants. Importantly, Mito-MES at nanomolar concentrations had additive antiviral and anti-inflammatory activity with DMF, an anti-inflammatory drug in multiple sclerosis (*11*). Finally, we showed that Mito-MES treatment can drastically reduce the replication of SARS-CoV-2 *in vivo* in a mouse model of SARS-CoV-2 infection.

To date, there is no safe, efficacious oral antiviral that is effective against SARS-CoV-2 variants, has *anti-inflammatory activity* and can *also* be given *long term* in humans. The favorable antiviral, antioxidant and anti-inflammatory properties of Mito-MES and its excellent safety profile in humans(*5*, *6*), can establish Mito-MES as a novel therapeutic strategy for outpatient treatment of mild to moderate acute COVID-19, for post-exposure prophylaxis against SARS-CoV-2 in high-risk exposures, for post-acute COVID-19 syndrome (PACS)(*1*) and as preexposure prophylaxis in high risk (un)vaccinated or immunocompromised patients where the SARS-CoV-2 vaccines may have low efficacy. Mito-MES is available as a dietary supplement and its safety for up to one year has been validated in clinical trials for oxidative damage-related diseases (*5*, *6*). Mito-MES may represent a rapidly applicable therapeutic strategy that can be added in the therapeutic arsenal against COVID-19.

## Author contributions

Conceptualization: TK

Methodology: AP, MS, MD, SS, HV, CH, ER, BJD, EF, GG, STR, CS, AP, BNG, GAF, VA, ML, OSH, TK

Investigation: AP, MS, MD, SS, HV, CH, ER, BJD, EF, GG, STR, CS, AP, BNG, GAF, VA, ML, OSH, TK

Visualization: AP, SS, TK

Funding acquisition: TK

Project administration: TK

Supervision: TK, OSH

Writing – original draft: AP, MS, TK

Writing – review & editing: AP, MS, SS, ML, OSH, TK

## Acknowledgments

The flow cytometry machine used in the study was purchased through the UCLA Center for AIDS Research (P30AI28697) grant. We thank Dani Dagan for helpful discussions and advice.

## STAR METHODS

### RESOURCE AVAILABILITY

#### Lead Contact

Further information and requests for resources and reagents should be directed to and will be fulfilled by the Lead Contact, Theodoros Kelesidis.

#### Materials availability

This study did not generate new unique reagents.

#### Data and code availability

All data are available in the main text or the supplementary materials.

### EXPERIMENTAL MODEL AND SUBJECT DETAILS

#### MATERIALS

##### Cells

Human adenocarcinoma lung epithelial Calu3 cells (Cat# HTB-55) and African green monkey kidney epithelial Vero-E6 cells (Cat# CRL-1586) were purchased from American Type Culture Collection (ATCC) (Manassas, VA). Human embryonic kidney 293T cells (HEK293T) cells stably expressing human angiotensin-converting enzyme 2 (h-ACE2 HEK293T cells) (Cat# SL221) were purchased from Genecopoeia (Rockville, MD). Murine fibroblast 17Cl-1 cells (Cat# NR-53719) and human lung carcinoma cells (A549) expressing human angiotensin-converting enzyme 2 (hACE2-A549) cells (NR-53821) were obtained through BEI Resources (Manassas, VA). Normal human bronchial epithelial cells (NHBE) (Cat# CC-2540) and Human primary small airway epithelial cells (HSAECs) (Cat# CC-2547) were obtained from Lonza (Basel, Switzerland), and all samples were de-identified. Lonza lung samples were obtained from donors ranging between 30-50 years and represented both males and females.

##### Viruses

The following reagents were obtained through Biodefense and Emerging Infectious (BEI) Resources of National Institute of Allergy and Infectious Diseases (NIAID), National Institutes of Health (NIH) (Manassas, VA): SARS-CoV-2, isolates 1) 2019-nCoV/USA-WA1/2020 strain, (GenBank accession no. MN985325.1) (labelled as wild type WT in experiments), 2) Isolate hCoV-19/USA/MD-HP01542/2021 (Lineage B.1.351, Beta Variant), NR-55282, contributed by Andrew S. Pekosz, 3) hCoV-19/USA/PHC658/2021 (Lineage B.1.617.2; Delta Variant), NR-55611, contributed by Dr. Richard Webby and Dr. Anami Patel, 4) Isolate hCoV-19/USA/MD-HP05285/2021 (Lineage B.1.617.2; Delta Variant), NR-55671, contributed by Andrew S. Pekosz, 5) Isolate hCoV-19/USA/GA-EHC-2811C/2021 (Lineage B.1.1.529; Omicron Variant), NR-56481, contributed by Mehul Suthar. 6)Recombinant Murine Coronavirus, icA59, NR-43000, contributed by Susan R. Weiss. The fluorescent reporter a stable mNeonGreen SARS-CoV-2 virus (icSARS-CoV-2-mNG) was gifted by Pei-Yong Shi’s lab at the University of Texas Medical Branch (UTMB). The Murine Hepatitis Virus A49 Green Fluorescent Protein (MHV-A49-GFP) was gifted by Volker Thiel’s lab at the University of Bern and Elke Mühlberger’s lab at Boston University.

##### Antibodies

Rabbit anti-SARS-CoV-2 nucleocapsid protein (NP) (clone ARC2372, Cat# PIMA536086) was purchased from Thermo Fisher Scientific (Waltham, MA). Rabbit anti-SARS-CoV-2 NP (polyclonal, Cat# 200401A50) was purchased from Rockland Immunochemicals (Pottstown, PA). Rabbit anti-SARS-CoV-2 (2019-nCoV) Spike S1 (clone 007, Cat# 40150-R007) was purchased from Sino Biologicals (Beijing, China). Mouse anti-SARS-CoV/SARS-CoV-2 (COVID-19) Spike (clone 1A9, Cat# GTX632604) was purchased from Genetex (Irvine, CA). Mouse anti-murine hepatitis virus (MHV) NP (clone 1.16.1, Cat# NR-45106) was obtained from BEI resources. Rabbit anti-human mitochondrial antiviral-signaling protein (MAVS) (clone D5A9E, Cat# 24930), rabbit anti-human/mouse Alexa Fluor 647 cleaved Caspase-3 (Asp175) (clone D3E9, Cat# 9602S), and rabbit anti-cleaved Caspase-3 (polyclonal, Cat# 9661) were purchased from Cell Signaling Technologies (Danvers, MA). Rabbit anti-human translocase of the mitochondrial outer membrane 70 (TOM70) (polyclonal, Cat# 14528-1-AP), anti-human/mouse myxovirus resistance protein 1 (MX1) (polyclonal, Cat# 13750-1-AP), and rabbit anti-human/mouse nuclear factor erythroid 2-related factor 2 (NRF2/NFE2L2) (polyclonal, Cat#16396-1-AP) were purchased from ProteinTech (Rosemont, IL). Mouse anti□human/rat heme oxygenase-1 (HMOX1/HO□1) (clone HO□1□2, Cat# LS□C343604) was purchased from LSBio (Seattle, WA).

The following secondary antibodies were purchased: goat anti-mouse Horseradish Peroxidase (HRP) (Cat# A16066), goat anti-rabbit HRP (Cat# PI31460), goat anti-mouse DyLight 488 (Cat# PI35503, goat anti-mouse DyLight 650 (Cat# PISA510174), goat anti-rabbit Alexa Fluor 488 (Cat# PI35553), goat anti-rabbit DyLight 650 (Cat# PISA510034) were purchased from Thermo Fisher Scientific.

##### Immunoassays

The following enzyme-linked immunosorbent assays (ELISA) were purchased: Legend Max Human IL-29 [interferon (IFN)-λ1)] ELISA Kit (Cat# 446307) was purchased from BioLegend (San Diego, CA). Anti-SARS-CoV-2 Nucleocapsid Protein Sandwich ELISA Kit (Cat# GTX535824) was purchased from Genetex (Irvine, CA). Human IFN-beta Quantikine ELISA kit (Cat# DIFNB0), Human interleukin (IL)-6 Quantikine ELISA kit (Cat# D6050) were purchased from Bio-Techne (Minneapolis, MN). The following multiplex Luminex immunoassays were purchased: Mouse ProcartaPlex kit [4-plex: interleukin-1β (IL-1β), IL-18, IL-6, and Tumor Necrosis Factor alpha (TNF-α)] was purchased from Thermo Fisher Scientific (Waltham, MA). Human High Sensitivity Cytokine A Premixed Luminex Performance Assay kit [6-plex: IL-1β, IL-6, IL-8, IL-10, TNF-α, and Vascular Endothelial Growth Factor (VEGF)] (Cat# FCSTM09-06) was purchased from Thermo Fisher Scientific (Waltham, MA).

##### Other reagents

The following materials were purchased from MilliporeSigma (Burlington, MA): Antimycin A (Cat# A8674-100MG), Bovine Serum Albumin (BSA) (Cat# A9576-50ML), Dimethyl fumarate (DMF) (Cat# 242926-25G), Dimethyl Sulfoxide (DMSO) (Cat# 472301-100ML), Donkey Serum (Cat# 566460-5ML), Goat Serum (Cat# NS02L-1ML), Ketamine hydrochloride/xylazine hydrochloride solution (Cat# K113-10ML), Mito-TEMPO (Cat# SML0737-5MG), Mix-n-Stain CF488 (Cat#MX488AS100), and Mix-n-Stain CF647 (Cat#MX647S100), Oligomycin (Cat# 495455-10MG), Rotenone (Cat# R8875-5G), Triton X-100 (Cat# T8787-50ML).

The following materials were purchased from Thermo Fisher Scientific (Waltham, MA): Corning™ Costar™ 96-Well, Cell Culture-Treated, Flat-Bottom Microplate (Cat#07-200-91), 6-diamidino-2-phenylindole (DAPI) (Cat# EN62248), Dihydroethidum (DHE)(Cat# D11347), Dulbecco’s modified eagle medium (DMEM) (Cat# 10-569-044), Dulbecco’s modified eagle medium (DMEM) low glucose (Cat# 11054-020), Epredia Gill™ Hematoxylin 3 (Cat# 22-050-203), Epredia™ Richard-Allan Scientific™ Eosin-Y with Phloxine (Cat#22-050-197), Epredia Richard-Allan Scientific Masson Trichrome Kit (Cat# 22-110-648), Halt Protease Inhibitor Cocktail (Cat# PI87785), Hanks buffered salt solution (HBSS) (Cat# 14-025-092), Hoechst 33342 (Cat# H3570), Hygromycin (Cat# 10-687-010), L-Glutamine 200 mM (Cat# 25-030-081), Methanol (Cat# AC423950040), MitoSOX Red (Cat# M36008), Eagle’s minimum essential medium (MEM) (Cat# MT10009CV), non-essential amino acids 100X (Cat# SH3023801), Pierce™ Bicinchoninic Acid Assay (BCA) protein assay kit (Cat# PI23227),Penicillin/Streptomycin 100X (Cat# 15-140-122), PowerUp^®^ SYBR Green RT-qPCR Master Mix (Cat# A25742), RevertAid first strand cDNA synthesis kit (Cat# FERK1622), Sodium Pyruvate 100 mM (Cat#11-360-070), sterile nylon 40 μm Filter (Cat# 07-201-430), 3,3’,5,5’-tetramethylbenzidine (TMB) peroxidase substrate (Cat# N301), Tissue-Plus™ optimal cutting temperature (O.C.T.) Compound (Cat# 23-730-571), T-PER™ tissue protein extraction reagent (Cat# PI78510), and trypsin-ethylenediamine tetraacetic acid (EDTA) (0.25% w/v).

The following reagents were purchased from Cayman Chemical (Ann Arbor, MI): Mitoquinone mesylate (Mito-MES) (Cat# 29317), Mitoquinol mesylate (Cat# 89950), Precellys 7 ml empty tubes (Cat#13917), Precellys 5.0 mm Zirconium Oxide beads (Cat# 13765), Precellys 2.8 mm Zirconium Oxide beads (Cat# 10401), Precellys 2 ml Soft Tissue Homogenizing Ceramic Beads Kit (CK14) (Cat# 10011152), and Remdesivir (Cat# 30354)

(1-Decyl) triphenylphoshonium bromide (DTPP) (Cat# sc-264801) was purchased from Santa Cruz Biotechnology (Dallas, TX). Brusatol (Cat# HY-19543) was purchased from MedChemExpress (Monmouth Junction, NJ). Fetal Bovine Serum (FBS) (Cat# 100-500) was purchased from GeminiBio (West Sacramento, CA). Costar 6.5 mm Transwell^®^ 0.4μm pore polyester membrane inserts (Cat# 38024), PneumaCult Ex-Plus media (Cat# 05040) and PneumaCult Air-Liquid Interface (ALI)-S media (Cat# 05050) were purchased from Stem Cell Technologies (Vancouver, Canada). XTT Cell Proliferation Assay Kit (Cat# 30-1011K) was purchased from ATCC. Ghost Dye Red 780 (Cat# 13-0865-T100) and Ghost Dye Violet 450 (Cat# 13-0863-T100) were purchased from Tonbo Biosciences (San Diego, CA). Direct-zol Ribonucleic acid (RNA) Miniprep Plus kit (Cat# R2072) was purchased from Zymo Research (Irvine, CA). Seahorse XF base media (Cat# 102353-100) and Seahorse XFp cell culture miniplate (Cat# 103025-100) were purchased from Agilent (Santa Clara, CA). Carbonylcyanide-4-(trifluoromethoxy)-phenylhydrazone (FCCP) (Cat# CM120-0010) was purchased from Enzo Life Sciences (Farmingdale, NY). Stop Solution 2N Sulfuric Acid (Cat# DY994) was purchased from Bio-Techne (Minneapolis, MN). Intracellular Staining Permeabilization Wash Buffer (10x) (Cat# 421002), Human TruStain FcX (Cat# 422302) and TruStain Fcx (anti-mouse CD16/32) (Cat#101320) were purchased from BioLegend (San Diego, CA). 32% w/v Paraformaldehyde (PFA) aqueous solution (Cat# 15714-1L) was purchased from Electron Microscopy Sciences (Hatfield, PA). Black 96 well μCLEAR® cell culture microplate (Cat# 655090) was purchased from Greiner Bio-One (Kremsmünster, Austria). Teklad Global 14% w/w protein rodent maintenance diet (Cat# 2914) was purchased from Envigo (Indianapolis, IN). RNeasy Mini Kit (Cat# 74104) was purchased from Qiagen (Hilden, Germany).

To study the antiviral and anti-inflammatory activity of the actual compound contained in the diet supplement given in humans (mitoquinol mesylate), the commercially available mitoquinol mesylate capsules were purchased for research purposes by MitoQ Ltd (New Zealand).

##### Mice

Tg(K18-ACE2)2Prlmn (Strain B6.Cg-Tg(K18-ACE2)2Prlmn/J (K18-hACE2) mice were purchased from the Jackson Laboratory (strain 034860).

#### METHODS

##### Cell cultures

Calu3, Vero-E6, HEK293-ACE2, Murine 17Cl-1 cells were maintained at 37 °C and 5% CO_2_ in DMEM or MEM supplemented with 10% (v/v) FBS, penicillin (100 units/ml), and streptomycin (100 μg/ml) (1X P/S). HEK293-ACE2 cells were maintained at 37 °C and 5% CO_2_ in MEM supplemented with 10% (v/v) FBS and hygromycin. h-ACE2 A549 cells were maintained at 37 °C and 5% CO_2_ in DMEM supplemented with 10% (v/v) FBS and blastidicin. Primary mouse lung cells that express human ACE2 were isolated from uninfected K18-hACE2 mice and were maintained at 37 °C and 5% CO_2_ in DMEM supplemented with 10% (v/v) FBS and 1X P/S.

##### Human Tissue Procurement

Large airways and bronchial tissues were acquired from de-identified normal human donors after lung transplantations at the Ronald Reagan University of California, Los Angeles (UCLA) Medical Center. Tissues were procured under Institutional Review Board-approved protocols at the David Geffen School of Medicine at UCLA. Human airway basal stem cells (ABSCs) from one biological replicate were used for two experiments. The biological replicates were from ABSCs isolated from lung transplant donors. No demographic data was available for the normal lung donor samples.

##### ABSC Isolation

Human airway basal stem cells (ABSCs) were isolated following a previously published method by our group(*18*). Briefly, airways were dissected, cleaned, and incubated in 16 units/ml dispase for 30 minutes (min) at room temperature. Tissues were then incubated in 0.5 mg/ml DNase for another 30 min at room temperature. Epithelium was stripped and incubated in 0.1% (v/v) Trypsin-EDTA for 30 min shaking at 37 °C to generate a single cell suspension. Isolated cells were passed through a 40 μm strainer and plated for Air-Liquid Interface cultures.

##### ABSC Air-Liquid Interface Cultures (Upper airway ALI cultures)

24-well 6.5 mm transwells with 0.4 μm pore polyester membrane inserts were coated with collagen type I dissolved in cell culture grade water at a ratio of 1:10. 100 μl was added to each transwell and allowed to air dry. ABSCs were seeded at 100,000 cells per well directly onto collagen-coated transwells and allowed to grow in the submerged phase of culture for 4-5 days with 500 μl media in the basal chamber and 200 μl media in the apical chamber. ALI cultures were then established and cultured with only 500 μl media in the basal chamber, and cultures were infected with SARS-COV-2 as indicated. Media was changed every other day and cultures were maintained at 37 °C and 5% CO_2_.

##### Human Primary Small Airway Epithelial Cells (HSAECs) Lower Respiratory Epithelium Airway ALI cultures

HSAECs were seeded onto collagen coated transwells and grown in the submerged phase of culture for 4–5 days in PneumaCult Ex Plus media with 500□μl media in the basal chamber and 200□μl media in the apical chamber. ALI cultures were then maintained for 21 days with only 500□μl PneumaCult ALI media in the basal chamber, and media changed every 2 days. Cultures were maintained at 37□°C and 5% CO_2_.

##### SARS-CoV-2 infection

All studies involving live virus were conducted at the UCLA Biosafety Level 3 (BSL3) high-containment facility with appropriate institutional biosafety approvals. SARS-CoV-2 was passaged once in Vero-E6 cells and viral stocks were aliquoted and stored at −80 °C. Virus titer was measured in Vero-E6 cells by median tissue culture infectious dose (TCID_50_) assay. Cell cultures in 96 well plates and ALI cultures were infected with SARS-CoV-2 viral inoculum [Multiplicity of infection (MOI) of 0.1 or at least 1 for ALI; 100 μl/well] prepared in media. For infection of Calu3 cells in 96 well plates with the fluorescent reporter a stable mNeonGreen SARS-CoV-2 virus (icSARS-CoV-2-mNG), an MOI of 0.3 and 100 μl/well was used. For infection of hACE2-A549 cells in 96 well plates an MOI of 0.5 or 1 and 50 μl/well was used. For mock infection, conditioned media (50 or 100 μl/well) alone was added.

##### Assessment of SARS-CoV-2 infection among different cell culture systems

SARS-CoV-2 infection was assessed by independent experiments that determined either the single cell content of SARS-CoV-2 or the total amount of secreted, intracellular or total (secreted and intracellular) SARS-CoV-2. The intracellular content of the SARS-CoV-2 nucleocapsid protein (NP) or the Spike (S) protein was assessed using ELISA, flow cytometry and immunofluorescence as described in Methods. Protein extract concentration was quantified using the Pierce BCA Protein Assay, according to manufacturer instructions (Thermo Fisher Scientific, Waltham, MA). SARS-CoV-2 NP or S protein content was expressed as either the percentage of cells that were positive for the SARS-CoV-2 viral protein (immunofluorescence or flow cytometry) or the median fluorescence intensity (MFI) of the SARS-CoV-2 viral protein per cell (immunofluorescence or flow cytometry). In ELISA experiments the intracellular SARS-CoV-2 NP or S protein content (pg) was normalized by the total cellular amount of protein within each experimental well (pg of viral protein per μg of total protein). The intracellular genomic expression of SARS-CoV-2 was assessed using primers specific for SARS-CoV-2 and real time PCR as described in Methods. The secreted amount of live SARS-CoV-2 in cell culture supernatant of infected cell cultures was assessed using viral titer as described in Methods. Percent infection was quantified as ((Infected cells/Total cells) - Background) *100 and the vehicle control was then set to 100% infection for analysis. The half maximal inhibitory concentration (IC_50_) for each experiment were determined using the Prism (GraphPad Holdings, San Diego, CA) software.

##### Viral titers

Infectious titers were quantified by limiting dilution titration using Vero-E6 cells. Briefly, Vero-E6 cells were seeded in 96-well plates at 5,000 or 10,000 cells/well. The next day, SARS-CoV-2-containing supernatant was applied at serial 10-fold dilutions ranging from 10^-1^ to 10^-8^ and, after 3-5 days, viral cytopathic effect (CPE) was assessed by microscopy or by determination of the intracellular SARS-CoV-2 NP using ELISA. TCID_50_/ml was calculated using the Reed-Muench method. For MHV-A59 Infectious titers were quantified by limiting dilution titration using Murine 17Cl-1 cells.

##### In-cell SARS-CoV-2 ELISA

To independently establish detection of SARS-CoV-2 infection using a more quantitative method to assess viral titer (not based on microscopy), we utilized in-cell SARS-CoV-2 ELISA based on intracellular detection of the SARS-CoV-2 S or NP protein. 5,000 or 10,000 Vero-E6 cells were seeded in 96 well plates in 100 μl. The next day, the cells were inoculated with 10 μl of a 10-fold titration series of SARS-CoV-2. Two to three days later, SARS-CoV-2 S or NP protein staining was assessed using an anti-SARS-CoV-2 S or NP protein antibody. Cells were fixed by adding 100 μl 8% (v/v) PFA to 100 μl of medium (final 4% solution) and 30 min of room temperature incubation. Medium was then discarded and the cells permeabilized for 5 min at room temperature by adding 100 μl of Intracellular Staining Permeabilization Wash Buffer (BioLegend). Cells were then washed with PBS and stained with 1:5,000 (anti-Spike S antibody clone 1A9) or 1:10,000 (anti-NP antibody clone ARC2372) in permeabilization buffer at 37 °C. After 1 hour, the cells were washed three times with washing buffer before a secondary anti-mouse or anti-rabbit antibody conjugated with HRP was added (1:20,000) and incubated for 1 hour at 37 °C. Following three times of washing, the 3,3’,5,5’-tetramethylbenzidine (TMB) peroxidase substrate was added. After 5 min light-protected incubation at room temperature, reaction was stopped using 0.5 M H2SO4. The optical density (OD) was recorded at 450 nm and baseline corrected for 620 nm using the Biotek microplate reader (Agilent Technologies, Santa Clara, CA).

##### Drug treatments

A concentration of Mito-MES between 10-1000 nM has been shown to be physiologically relevant, efficacious and non-cytotoxic in human mammalian cells. The antiviral activity of Mito-MES was evaluated in Calu3, Vero-E6, HEK293T, h-ACE2 A549 and HSAECs ALI cell cultures. All SARS-CoV-2 studies were performed in biological triplicate. Cultured cells were incubated separately with Mito-MES or decyl triphenyl phosphonium cation (DTPP)— a non-antioxidant mitochondria-targeted control compound which allowed for the effects of the TPP component of Mito-MES to be examined. All drug concentrations were between 10 nM-100 μM as shown in the figures. The concentration of DMSO vehicle control was maintained constant at 0.1% v/v for all treatments. Drug effects were measured relative to vehicle controls *in vitro*. Unless stated, Calu3, Vero-E6, HEK293-ACE2, and ALI cultures were pretreated for at least 1□hour and up to 24 hours (Brusatol, DMF) with the indicated treatments (Mito-MES, DMF, Brusatol, Mito-TEMPO) or vehicle control. The cells were then washed, infected with SARS-CoV-2 for 2□hrs, the virus was removed, and the treatments (Mito-MES, mito-TEMPO, remdesivir) were added back. Remdesivir, well characterized direct acting antiviral agent, was used as antiviral control.

##### Assessment of cell cytotoxicity

To measure cell viability to determine if there was any treatment-induced cytotoxicity, uninfected cells were plated and treated with the same compound dilutions used for the *in vitro* efficacy studies. As above, 0.01% DMSO-treated cells served as the 0% cytotoxicity control. After 24-48 hrs, cell viability was measured on a Synergy 2 Biotek microplate reader (Agilent Technologies, Santa Clara, CA) via the XTT Cell Proliferation Assay Kit according to the manufacturer’s protocol (ATCC, Manassas, VA). Similar data were obtained in three independent experiments. LDH cytotoxicity assay was also used to assess cytotoxicity in cell culture supernatant according to the manufacturer’s protocol (Sigma-Aldrich). Flow cytometry was used to independently assess cell viability and levels of target proteins in single viable cells.

##### Imaging and immunofluorescence

After 24-48 hrs of SARS-CoV-2 infection (or mock), live cell images were obtained by wide field fluorescence microscopy using a 10x air objective (Leica DM IRB, Wetzlar, Germany). For immunofluorescence, separate wells of cells were fixed with 4% paraformaldehyde in phosphate-buffered saline (PBS) for 20 min. The fixed samples were then permeabilized and blocked for 1 hour in a “blocking solution” containing PBS with 2% bovine serum albumin, 5% donkey serum, 5% goat serum, and 0.3% Triton X-100. Primary antibodies were diluted in the blocking solution and added to samples overnight at 4 °C. The following antibodies and dilutions were used: polyclonal rabbit anti-Tom70 (1:100), rabbit anti-SARS-CoV-2 Spike S1 (clone #007) (1:100), polyclonal rabbit anti-cleaved caspase-3 (1:200), polyclonal rabbit anti-SARS-CoV-2 NP (1:1000). Samples were then rinsed 5 times for 2 min each with PBS containing 0.3% (v/v) Triton X-100, followed by incubation with fluorescent-conjugated secondary antibodies diluted 1:1,000 in blocking buffer for 2 hrs at room temperature. Secondary antibodies were goat anti-rabbit Alexa Fluor 488 IgG, goat anti-mouse Alexa Fluor 546, goat anti-rabbit DyLight 650. Samples were then rinsed 5 times for 2 min each with PBS containing 0.3% Triton X-100, followed by DAPI diluted in PBS at 1:5000 for 10 min. Immunofluorescence (IF) images were obtained using an LSM880 Zeiss confocal microscope (Carl Zeiss GmbH, Jena, Germany) with Airyscan using a 20X air objective or a 63X Apochromat oil-immersion objective. Immunofluorescence images were quantified using CellProfiler 2.0(*19*) (Broad Institute, Cambridge, MA). DAPI was used to count total cell numbers in order to obtain a percentage of cells positive for spike protein, cleaved caspase-3, or TOM70. Alternatively, 96-well plates were imaged using the Operetta (PerkinElmer, Waltham, MA) system.

##### High-throughput imaging

Cells plated in Greiner μClear 96-well plates were stained with antibodies for immunofluorescence as above and were imaged on Operetta high-content wide-field fluorescence imaging system using a 10X air objective. Fifteen fields were imaged per well with a similar field distribution across all wells. CellProfiler 2.0 was used to quantify cell counts and infection ratios(*19*).

##### Cellular reactive oxygen species analysis

Vero-E6 or Calu3 cells were treated and infected with icSARS-CoV-2-mNG reporter virus as indicated for 24-48 hrs. Subsequently, cells were washed once with hanks buffered salt solution (HBSS) with calcium and magnesium, without phenol-red and stained for 10 min with 1 μg/ml Hoechst 33342 and 5 μM dihydroethidum (DHE) in HBSS. Following two washes with HBSS, cells were returned to an imaging medium without phenol red (low glucose DMEM, supplemented with 10% (v/v) FBS, 2 mM glutamine and 1 mM sodium pyruvate and 1X P/S and imaged with a Leica DM IRB wide field fluorescence microscope as described above using DAPI, GFP, and RFP filters. CellProfiler 2.0 was used to quantify cell counts, infection ratios per well, and DHE fluorescence intensity per cell(*19*).

##### Flow cytometry

Within the UCLA BSL3 high-containment facility, SARS-CoV-2 infected and uninfected cells were resuspended in PBS and single cell suspensions were incubated with viability dye at 1:500 (Fixable Ghost Dye Red 780 or Ghost Dye Violet 450) for 20 min in the dark at room temperature. Cells were washed and appropriate antibodies were added to each tube and incubated in the dark for 20 min on ice. Cells were then washed and fixed with 4% (v/v) paraformaldehyde or methanol for 30 min at 4 °C. Cells were then washed and were transferred in polypropylene Eppendorf tubes (E-tubes) to BSL2 containment facility for further processing. Appropriate antibodies were added to each tube and incubated in the dark for 20 min on ice.

The following antibodies were used for staining in flow cytometry: rabbit anti-SARS-CoV-2 NP (clone ARC2372. 1:50), rabbit anti-SARS-CoV-2 Spike (clone 007, 1:50), mouse anti-SARS-CoV-2 Spike (clone 1A9, 1:50), rabbit anti-human MAVS (clone D5A9E, 1:10), rabbit anti-human TOM70 (polyclonal,1:5), rabbit anti-human/mouse Alexa Fluor 647 cleaved caspase-3 (clone D3E9, 1:25), mouse anti□human, HO□1 (clone HO□1□2, 1:20), anti-MX1 (polyclonal, 1:10), rabbit anti-NRF2 (polyclonal, 1:50). Non-fluorescent unconjugated primary antibodies were conjugated with Mix-n-Stain CF488 or CF647 Antibody Labeling Kits from MilliporeSigma (Burlington, MA).

The cells were washed twice with PBS and were transferred to tubes for fluorescence activated cell sorting (FACS) analysis. Samples were acquired using an LSRFortessa™ flow cytometer and FACSDiva™ software (Becton Dickinson, Franklin Lakes, NJ). Data were analyzed using FlowJo™ software (Becton Dickinson, Franklin Lakes, NJ). At least 10,000 cells were acquired for each analysis and only live and singlet cells were chosen for analysis and gating (i.e., dead cells and aggregates were excluded). Each representative flow plot was repeated at least 3 times. Single stain and also fluorescence minus one (FMO) controls were used in the presence of a given concentration of the antibody staining cocktail.

##### Mitochondrial Reactive oxygen species analysis

After 24 hrs post infection, cells were collected and stained with 5 μM MitoSOX Red Mitochondrial reactive oxygen species (mito-ROS) indicator for 30 min at 37 °C. Cells were washed with PBS, fixed with 4% paraformaldehyde for 30 min at 4 °C and transferred to polypropylene FACS tubes. Cells were then analyzed using an LSR Fortessa flow cytometer and FACSDiva software), and data were analyzed using FlowJo software (Becton Dickinson, Franklin Lakes, NJ).

##### Biomarkers of inflammation

Protein levels of secreted IL-6 were determined in cell culture supernatants using ELISA kits according to the manufacturer (Bio-Techne, Minneapolis, MN). Luminex immunoassay was used to measure human cytokines [interleukin-1β (IL-1β), IL-8, IL-10, TNF-α, Vascular endothelial growth factor (VEGF)] secreted by Calu3 cells and airway lung epithelial cells in cell culture supernatants according to the manufacturer (Bio-Techne, Minneapolis, MN). A 4-Plex ProcartaPlex assay (Thermo Fisher Scientific, Waltham, MA) was used to measure murine cytokines (IL-1β, IL-6, IL-18, TNF-α) in mouse lung protein lysates.

##### RNA extraction and Real-Time Quantitative Reverse Transcription Polymerase Chain Reaction (RT-qPCR)

Total RNA was isolated using the RNeasy Mini Kit or Direct-zol RNA Miniprep kit (Zymo Research) and complementary deoxyribonucleic acid (cDNA) was synthesized using oligo deoxythymine (dT) primers and RevertAid first strand cDNA synthesis kit. Quantitative real-time reverse transcription PCR was performed using SYBR Green Master Mix and primers specific for SARS-CoV-2 as well as glyceraldehyde 3-phosphate dehydrogenase (GAPDH) transcripts. The following primers were used: 2019-nCoV_N1-F: GACCCCAAAATCAGCGAAAT, 2019-nCoV_N1-R: TCTGGTTACTGCCAGTTGAATCTG; h-GAPDH-F: CCACCTTTGACGCTGGG; h-GAPDH-R: CATACCAGGAAATGAGCTTGACA. All qRT-PCR reactions were performed using BIO-RAD CFX96 Touch Real-Time PCR Detection System (Bio-Rad Laboratories, Hercules, CA) on 96-well plates. PCR reactions included SYBR Green RT-PCR Master Mix, 10 μM primers and 5 μl of cDNA. Reactions were incubated at 45 °C for 10 min for reverse transcription, 95 °C for 2 min, followed by 40 cycles of 95 °C for 15 seconds (sec) and 60 °C for 60 sec. Gene expression fold change was calculated with the Delta-delta-cycle threshold (DDCt) method. Viral RNA levels were depicted as fold change over mock infected samples.

##### Mitochondrial respirometry analysis

Calu3 cells were seeded at a density of 16,000 cells per well into 8-well Seahorse mini plates and allowed to attach for 24 hrs. Treatments were started 24 hrs before infection with SARS-CoV-2 (WA1 strain, MOI 0.1-0.5). Cells were infected for 48 hrs and subsequently analyzed by Seahorse respirometry. Briefly, growth media was replaced with Seahorse XF base media (supplemented with 2 mM glutamine and 1 mM sodium pyruvate, pH adjusted to 7.4) twice and respirometry assessed using a Seahorse XF HS Mini Analyzer (Agilent Santa Clara, CA) and the using the Mito Stress Test protocol. After basal respiration, mitochondrial proton leak was assessed after oligomycin injection, maximal respiration was assessed after Carbonylcyanide-4-(trifluoromethoxy)-phenylhydrazone (FCCP) injection, and non-mitochondrial respiration after rotenone, and antimycin A injection. Following the assay, cells were fixed in 4% (v/v) paraformaldehyde as described above and SARS-CoV-2 positive cells stained against SARS-CoV-2 nucleocapsid protein, and nuclei stained with DAPI. Wells were imaged using an LSM880 Zeiss confocal microscope (Carl Zeiss GmbH, Jena, Germany) with a 10X air objective and cell counts and infection ratios determined using CellProfiler 2.0. Agilent Wave 2.6 software (Agilent Santa Clara, CA) was used to normalize oxygen consumption rates to cell counts.

##### Mouse studies of SARS-CoV-2 infection

All the antiviral studies were performed in animal biosafety level 3 (BSL3) facility at University of California, Los Angeles. All work was conducted under protocols approved by the Institutional Animal Care and Use Committee (IACUC). We used a model of mice hemizygous for the expression of hACE2 gene (K18-ACE2) and we performed three studies in these mice to evaluate the *in vivo* efficacy of Mito-MES as antiviral and anti-inflammatory therapeutic agent in SARS-CoV-2 infection. In total, we utilized 60 male and female 4 to12-week-old specific pathogen–free hemizygous for Tg(K18-ACE2)2Prlmn (Strain B6.Cg-Tg(K18-ACE2)2Prlmn/J, the Jackson laboratory strain 034860) mice. Animals were housed in individually ventilated cages, 5 mice per cage, on a 12-h light–dark cycle at 21–23 °C and 40–60% humidity. Mice were allowed free access to irradiated standard rodent diet (Tecklad 2914C) and sterilized water.

Cohort A included 15 male K18-hACE2 mice between 4 to 8 weeks of age (16-25 g). Mice were treated intraperitoneally with either vehicle control (normal saline 10% DMSO; Ctrl) (n=10) or Mito-MES 4 mg/kg/day (n=5) overnight before the infection (dose 1). Ten mice were then given a second dose of vehicle or Mito-MES and after 2 hrs were infected intranasally with wild type (WT) SARS-CoV-2 [10,000 plaque-forming unit (PFU)/mouse in <50 μl of PBS, day 0]. Mice were anesthetized with a mixture of ketamine/xylazine before each intranasal infection. Five mice were given intraperitoneally Mito-MES (first dose) 8 hrs post infection (hpi)(n=5). Additional doses of Mito-MES (n=10) or vehicle (n=5) were given on day 1 (24 hpi), day 2 (48 hpi) and day 3 (72 hpi). The body weight of mice was measured each day. Three days post infection (dpi) animals were humanely euthanized. Whole left lungs were harvested and were processed to create homogenates for viral titration via TCID50, single cell suspension for flow cytometry, or protein lysates for immunoassays. The total (secreted and intracellular) amount of SARS-CoV-2 NP in lung tissue homogenates (protein lysates) from mice infected with SARS-CoV-2 was assessed by SARS-CoV-2 NP ELISA. The total (secreted and intracellular) amount of live SARS-CoV-2 in lung tissue homogenates from mice infected with SARS-CoV-2 was assessed by viral titer.

Cohort B included 20 female K18-hACE2 mice between 4 to 8 weeks of age (16-21 g). Mice were treated intraperitoneally with either vehicle control (normal saline 10% DMSO; Ctrl) (n=10) or Mito-MES 4 mg/kg/day (n=10) overnight before the infection (dose 1). Next day (Day 0), mice were then given a second dose of vehicle or Mito-MES and after 2 hrs were infected intranasally with Beta Variant of SARS-CoV-2 (10,000 PFU/mouse in <50 μl of PBS). Mice were anesthetized with a mixture of ketamine/xylazine before each intranasal infection. Additional doses of Mito-MES (n=10) or vehicle (n=10) were given on each day for up to 5-7 dpi. The body weight of mice was measured each day. 5-7 dpi animals were humanely euthanized. Whole left lungs were harvested and were processed for histopathology studies, single cell suspension for flow cytometry or protein lysates for immunoassays.

Cohort C included 20 male K18-hACE2 mice between 4 to 8 weeks of age (16-25 g). Mice were treated via gavage with either vehicle control (normal saline 10% DMSO; Ctrl) (n=10) or Mito-MES 20 mg/kg/day (n=10) overnight before the infection (dose 1). As source of Mito-MES the actual diet supplement capsule (5 mg Mito-MES per capsule) that is given in humans was utilized. Each capsule was opened and was dissolved in saline 10% DMSO solution before given via gavage to mice. Next day (Day 0), mice were then given a second dose of vehicle or Mito-MES and after 2 hrs were infected intranasally with Delta Variant of SARS-CoV-2 (10,000 PFU/mouse in <50 μl of PBS). Mice were anesthetized with a mixture of ketamine/xylazine before each intranasal infection. Additional doses of Mito-MES (n=10) or vehicle (n=10) were given on day 1, day 2 and day 3 post infection. The body weight of mice was measured each day. On 3 dpi animals were humanely euthanized. Whole left lungs were harvested and were processed for single cell suspension for flow cytometry or viral titration via TCID_50_.

##### Mouse tissue processing

Tissue samples were collected at necropsy. Less than 20 mg of lung tissue was either embedded in paraffin for hematoxylin and eosin (H&E) staining or was cryoembedded in Tissue-Plus O.C.T Compound for immunofluorescence staining. The right middle lobe was used for cell isolation and flow cytometry analysis. Tissues were weighted and <0.25 gram tissue samples were rinsed with DBPS, cut in 0.5 cm size pieces that were placed in 7 ml screwcap dissociation tubes with ceramic beads (Precellys) filled with 2.5 ml prewarmed (37 °C) digestion medium (2 mg/ml Collagenase, DNAse 0.1 mg/ml, 1% FCS). Tissues were mechanically dissociated at power 3,000 rpm for one 10-second cycle using a Precellys 24 homogenizer (Bertin Technologies, Montigny-le-Bretonneux, France), followed by a 30-minute incubation at 37 °C and another 10-second dissociation cycle (3,000 rpm). The homogenate was filtered through a sterile 40 μm nylon filter and cells were then processed for flow cytometry. The entire left lung was processed immediately for viral titer. Lung tissues were placed in 2 ml screwcap dissociation tubes with 60 × 1.4 mm ceramic beads at power 5,000 rpm using a Precellys 24 homogenizer and were mechanically dissociated in 0.5 ml DPBS for 90 sec. Homogenates were centrifuged at 16,000 g for 10 min and supernatants were then cryopreserved at −80 °C for viral titer in Vero-E6 cells. For tissue lysates that were used for protein measurements, 50-100 mg of lung tissue samples were placed in 2 ml screwcap dissociation tubes with ceramic beads (Precellys) and were mechanically dissociated in T-PER tissue protein extraction Reagent at power 5,000 rpm leaving samples to cool on ice between 1–2 repeated 20-second cycles as previously described(*20*).

##### Isolation of primary mouse lung cells

Following euthanasia, uninfected K18-hACE2 mice of identical age and gender like the mice in cohort C, were perfused with normal saline before removal of the left lung. Lung tissues were then rinsed with DBPS and processed similarly to SARS-CoV-2 infected mice to create cell suspension. Cells were then seeded in T75 flasks with complete medium [DMEM supplemented with 10% (v/v) FBS, 1X P/S] and were allowed to adhere overnight. The next day the medium was exchanged, and cells were maintained in complete medium before seeding in 96 well plates for SARS-CoV-2 infection and treatment with Mito-MES.

##### Mouse lung histological analysis

Paraffin-embedded murine lung tissue blocks were cut into 5 μm sections that were stained with hematoxylin and eosin (H&E) or Trichrome Masson. Light microscopic scans of whole lung were examined by an experienced pathologist using an Olympus BX53 microscope. Perivascular inflammation, airway inflammation, and alveolar inflammation were individually assessed semiquantitatively using a 5-point grading scheme considering the focality and intensity of the inflammatory infiltrate: 0 – absent, 1 – focal mild or patchy minimal, 2 – multifocal mild or focal moderate, 3 – diffuse moderate or focal severe, 4 – diffuse severe or with epithelial injury/necrosis. The overall grade of inflammation was determined using the sum of these three scores (max 12). Interstitial and airway fibrosis were absent in all cases, confirmed by Masson trichrome stain.

##### Statistics

Unless noted, error bars in all figures represent mean and standard error of means (SEM). In the figures, p-values are presented for comparisons between treatment groups and controls and are denoted by asterisks. To pool cells from different experiments, each measurement was first normalized to the vehicle controls of each experiment. Each experiment contains at least 3 biological replicates (number of wells) and each analysis contains at least 2 independent experiments. For flow analysis of cells, at least 10,000 events were acquired for the population of interest. For analysis of data that contains more than 2 groups, the Kruskal-Wallis test was performed to compare samples; if these comparisons had a p value less than 0.05 then Mann-Whitney U tests were used to compare statistical difference between 2 groups. P-values less than 0.05 by Kruskal-Wallis or Mann-Whitney were considered significant. Consultation on statistical analysis was performed with the UCLA Biostatistics Department. All analyses were performed with GraphPad, version 8.0 (GraphPad Holdings, San Diego, CA).

## References

1. A. Nalbandian et al., Post-acute COVID-19 syndrome. Nat Med 27, 601–615 (2021).

2. S. Zhou et al., beta-d-N4-hydroxycytidine Inhibits SARS-CoV-2 Through Lethal Mutagenesis But Is Also Mutagenic To Mammalian Cells. J Infect Dis 224, 415–419 (2021).

3. M. Hu et al., Respiratory syncytial virus co-opts host mitochondrial function to favour infectious virus production. Elife 8, (2019).

4. A. C. Codo et al., Elevated Glucose Levels Favor SARS-CoV-2 Infection and Monocyte Response through a HIF-1alpha/Glycolysis-Dependent Axis. Cell Metab 32, 437–446 e435 (2020).

5. R. A. Smith, M. P. Murphy, Animal and human studies with the mitochondria-targeted antioxidant MitoQ. Ann N Y Acad Sci 1201, 96–103 (2010).

6. M. J. Rossman et al., Chronic Supplementation With a Mitochondrial Antioxidant (MitoQ) Improves Vascular Function in Healthy Older Adults. Hypertension 71, 1056–1063 (2018).

7. J. Zielonka et al., Mitochondria-Targeted Triphenylphosphonium-Based Compounds: Syntheses, Mechanisms of Action, and Therapeutic and Diagnostic Applications. Chem Rev 117, 10043–10120 (2017).

8. B. D. Fink et al., Bioenergetic effects of mitochondrial-targeted coenzyme Q analogs in endothelial cells. J Pharmacol Exp Ther 342, 709–719 (2012).

9. S. Zhang, Q. Zhou, Y. Li, Y. Zhang, Y. Wu, MitoQ Modulates Lipopolysaccharide-Induced Intestinal Barrier Dysfunction via Regulating Nrf2 Signaling. Mediators Inflamm 2020, 3276148 (2020).

10. N. E. Saidu et al., Dimethyl Fumarate Controls the NRF2/DJ-1 Axis in Cancer Cells: Therapeutic Applications. Mol Cancer Ther 16, 529–539 (2017).

11. S. Saha, B. Buttari, E. Panieri, E. Profumo, L. Saso, An Overview of Nrf2 Signaling Pathway and Its Role in Inflammation. Molecules 25, (2020).

12. D. Olagnier et al., SARS-CoV2-mediated suppression of NRF2-signaling reveals potent antiviral and anti-inflammatory activity of 4-octyl-itaconate and dimethyl fumarate. Nat Commun 11, 4938 (2020).

13. X. Y. Liu, B. Wei, H. X. Shi, Y. F. Shan, C. Wang, Tom70 mediates activation of interferon regulatory factor 3 on mitochondria. Cell Res 20, 994–1011 (2010).

14. L. A. Sena, N. S. Chandel, Physiological roles of mitochondrial reactive oxygen species. Mol Cell 48, 158–167 (2012).

15. J. Bizzotto et al., SARS-CoV-2 Infection Boosts MX1 Antiviral Effector in COVID-19 Patients. iScience 23, 101585 (2020).

16. H. Cao et al., The anti-viral dynamin family member MxB participates in mitochondrial integrity. Nat Commun 11, 1048 (2020).

17. K. M. White et al., Plitidepsin has potent preclinical efficacy against SARS-CoV-2 by targeting the host protein eEF1A. Science 371, 926–931 (2021).

18. A. Purkayastha et al., Direct Exposure to SARS-CoV-2 and Cigarette Smoke Increases Infection Severity and Alters the Stem Cell-Derived Airway Repair Response. Cell Stem Cell 27, 869–875 e864 (2020).

19. L. Kamentsky et al., Improved structure, function and compatibility for CellProfiler: modular high-throughput image analysis software. Bioinformatics 27, 1179–1180 (2011).

20. M. Daskou et al., ApoA-I mimetics favorably impact cyclooxygenase 2 and bioactive lipids that may contribute to cardiometabolic syndrome in chronic treated HIV. Metabolism 124, 154888 (2021).

