## Supplemental Material for "Mitoquinone mesylate targets SARS-CoV-2 infection in preclinical models"

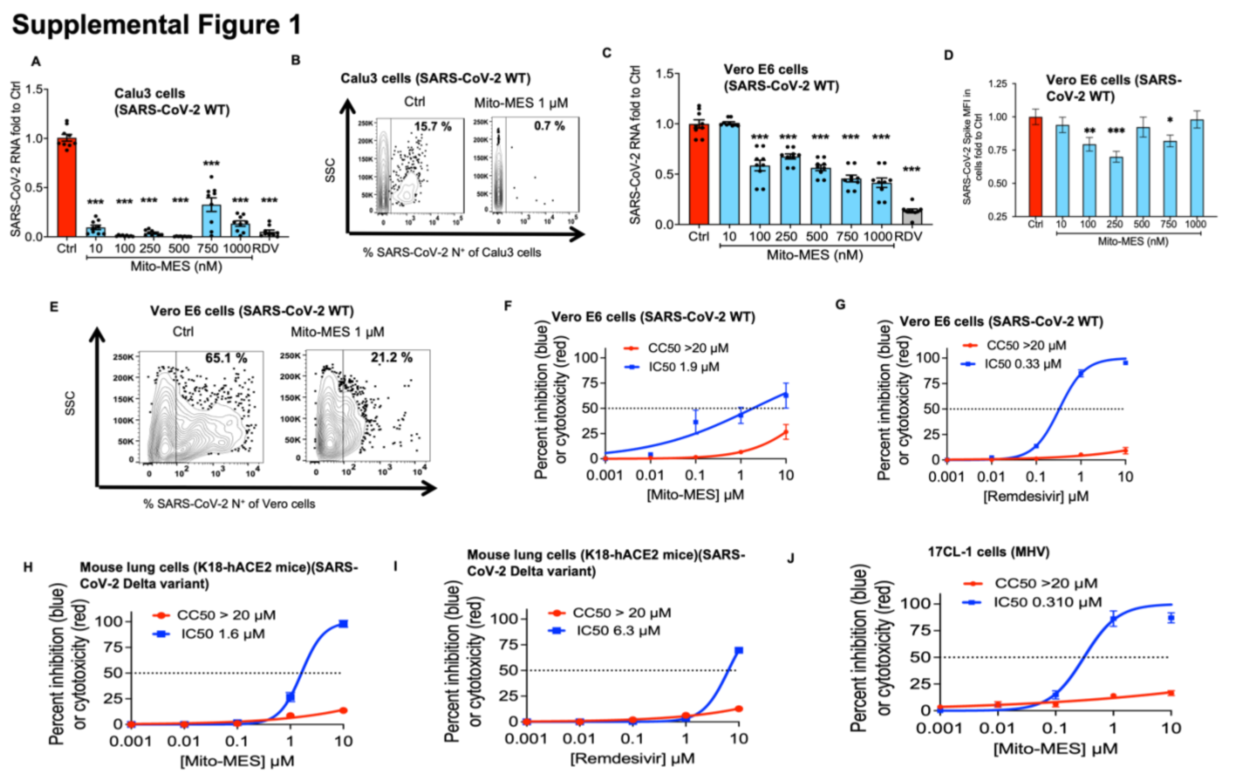


**Fig. S1. Mitoquinone mesylate (Mito-MES) has stronger anti-SARS-CoV-2 activity in human epithelial compared to non-human cell types.** (**A** to **J**). Vero E6, Calu3, 17CL-1 cells and lung cells isolated from K18-hACE2 mice were infected with wild type (WT) SARS-CoV-2, Beta, Delta variants or mouse hepatitis virus (MHV) at an MOI of 0.1 (48 hrs) and were treated with vehicle control (Ctrl) or drugs [Mito-MES or remdesivir (RDV)] for at least 1 hour before infection and throughout the experiment as shown. Viral replication at 48 hours post infection (hpi) by qPCR [(**A**), (**C**)], immunofluorescence (**D**), flow cytometry [(**B**)(**E**)] or TCID50-assay. Cell cytotoxicity was assessed in uninfected cells by the XTT (48 h) or the LDH assay (mouse cells). IC50, 50% cytotoxic concentration (CC50) values are indicated. In all panels summary data are presented as mean ± SEM or representative images of three or more experiments (with at least 2 replicates). Each data-point represents one biological sample. Unless otherwise stated, statistical comparison was done between the Ctrl and each shown experimental group by using two-tailed Mann–Whitney (*p < 0.05, **p < 0.01, ***p < 0.001).

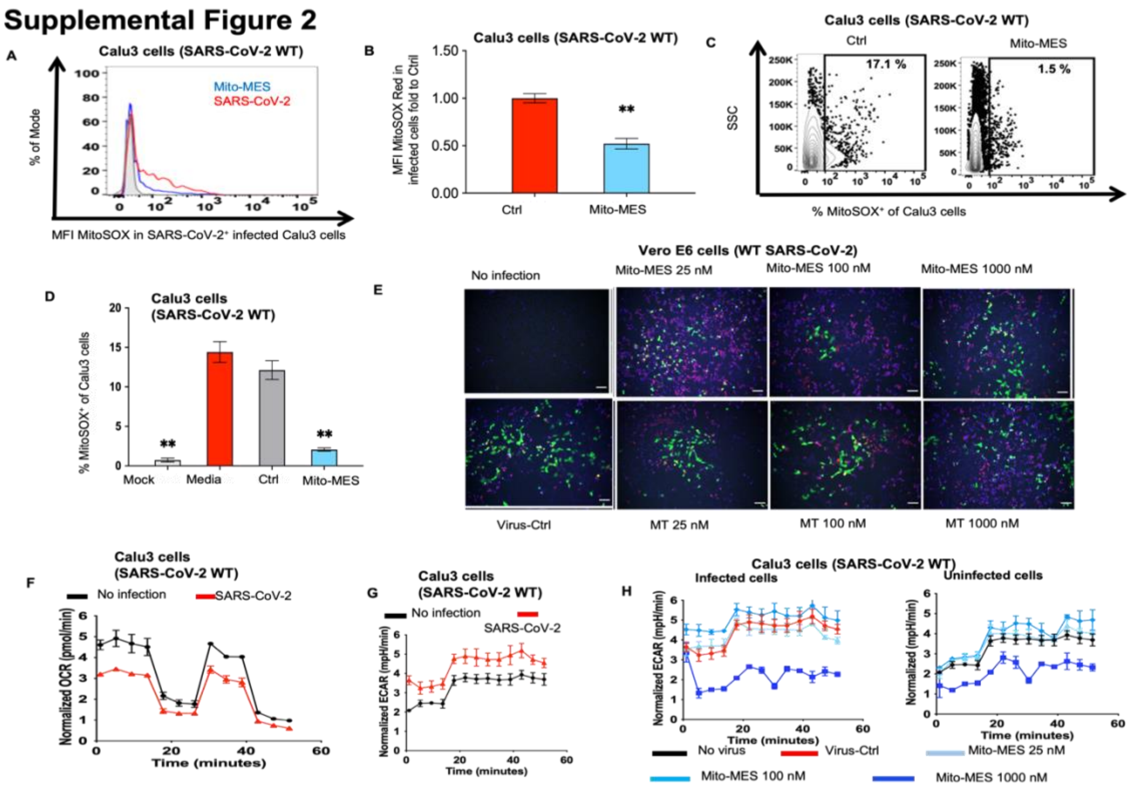


**Fig. S2. The antiviral activity of Mito-MES against SARS-CoV-2 in epithelial cells depends partially on its antioxidant activity.** (**A** to **H**) Calu3 cells and Vero-E6 cells (**E**) were infected with wild type (WT) SARS-CoV-2 at an MOI of 0.1 (48 hrs) or with fluorescent SARS-CoV-2 (MOI 0.3) (**E**) and were treated with vehicle control (Ctrl) or drugs [Mito-MES (100 nM unless indicated otherwise) or Mito-TEMPO (MT)] for at least 1 hour before infection and throughout the experiment as shown. Assessment of fluorescence [fluorescence intensity (MFI) (**A**), (**B**) and percent of positive cells (**C**), (**D**)] of MitoSOX Red in Calu3 cells. (**E**), Immunofluorescent analysis of viral replication and cellular oxidative stress [fluorescence of dihydroethidium (DHE)] at 48 hpi. (**F** to **H**) Seahorse XF Analyzer determined cellular bioenergetics [oxygen consumption rate (OCR) (**F**)(**G**) and extracellular acidification rate (ECAR) (**H**) in live (un)infected Calu3 cells treated with Mito-MES or Ctrl and infected with SARS-CoV-2. In all panels, data are representative or mean ± SEM of at least three experiments with at least 2 replicates. Statistical comparison was done between the Ctrl and each shown experimental group by using two-tailed Mann–Whitney (*$p < 0.05$, **$p < 0.01$, ***$p < 0.001$).

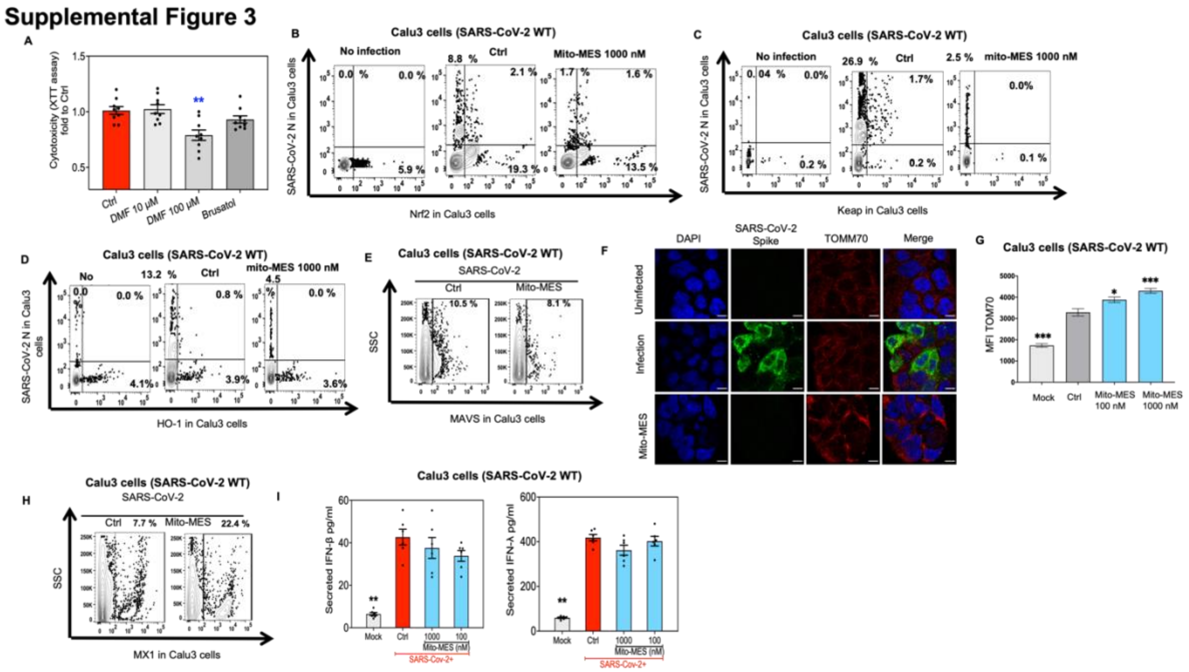


**Fig. S3. Impact of Mito-MES on host proteins that mediate cellular responses to SARS-CoV-2 infection in epithelial cells.** (**A**) Cytotoxicity by XTT assay in uninfected Calu3 cells treated as shown. (**B** to **I**) Calu3 cells were infected with wild type (WT) SARS-CoV-2 (MOI 0.1) and were treated with vehicle (Ctrl) or Mito-MES (1000 nM unless otherwise indicated) for at least 1 hour before infection and until 48 hours post infection (hpi) as shown. (**B** to **E** and **H**) Flow cytometric assessment of levels of key proteins of the Nrf2 pathway [Nrf2, Keap, Heme Oxygenase 1 (HO-1)], MAVS (**E**), MX1 (**H**) in (un)infected Calu3 cells. (**F**) Representative panels show immunostaining for SARS-CoV-2 spike protein (green) and TOM70 (red). DAPI stained cell nuclei. Scale bar= 100 μm. (**G**) Summary data of (**F**). (**I**) ELISA measured secreted interferons (beta: IFN-β, lambda: IFN-λ) in supernatants of (un)infected Calu3 cells (48 h). Data are representative for or pooled from at least three experiments with at least 2 replicates. Bars indicate mean ± SEM. Statistical comparison was done between the Ctrl and each shown experimental group by using two-tailed Mann–Whitney (*$p < 0.05$, **$p < 0.01$, ***$p < 0.001$).

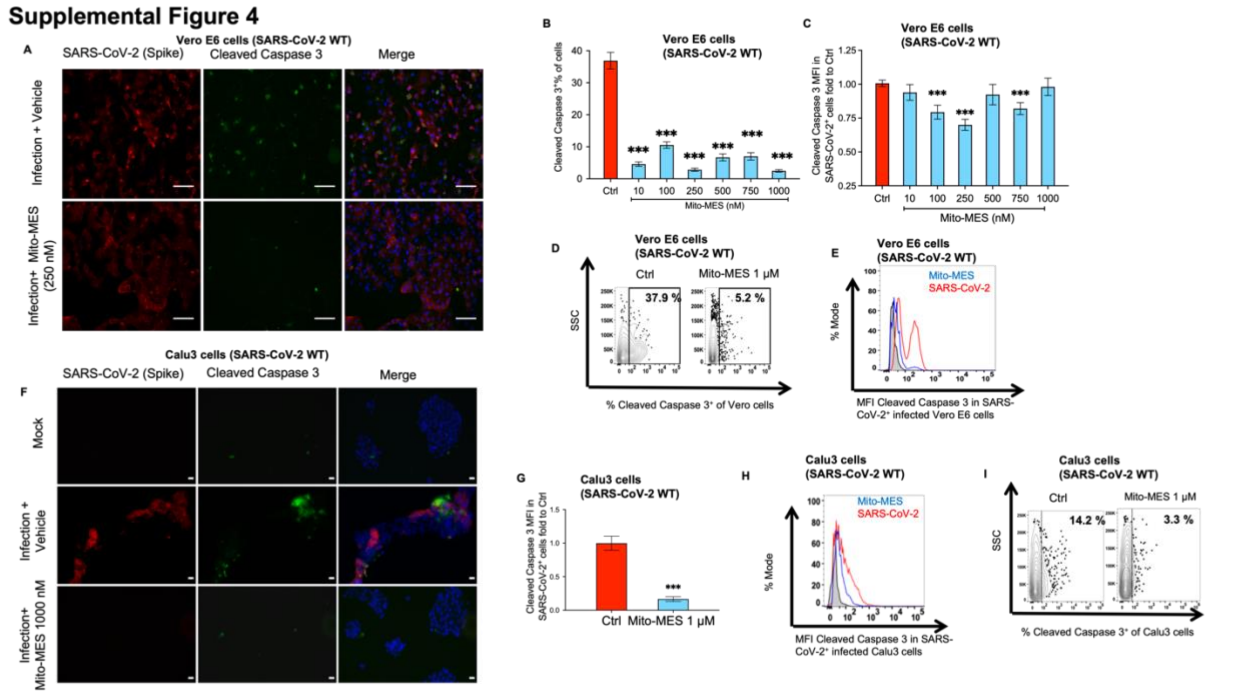


**Fig. S4. Mitoquinone mesylate restricts SARS-CoV-2 associated apoptosis in epithelial cells. (A** to **I)** Vero-E6 (**A** to **E**) and Calu3 (**F** to **I**) cells were infected with SARS-CoV-2 for 48 hrs (MOI of 0.1) and were treated with Mito-MES versus DMSO control (Ctrl) as shown. [(**A**), (**F**)] Immunostaining for SARS-CoV-2 spike protein (red) and apoptosis marker cleaved caspase-3 (green) at 48 hrs post-infection (hpi). DAPI stained cell nuclei. Scale bar=100 μm. [(**B**), (**C**), (**G**)] Summary of immunofluorescence data for protein levels [% of cells positive for protein or median fluorescence intensity (MFI)] of cleaved caspase-3 in Vero-E6 and Calu3 cells. Protein levels of cleaved caspase-3 were also assessed by flow cytometry [(**D**), (**E**), (**H**), (**I**)]. In all panels, data are shown as summary data (mean ± SEM) or representative images of three or more independent experiments with at least 2 replicates. Unless otherwise stated, statistical comparison was done between the Ctrl and each shown experimental group by using two-tailed Mann–Whitney (*p < 0.05, **p < 0.01, ***p < 0.001).

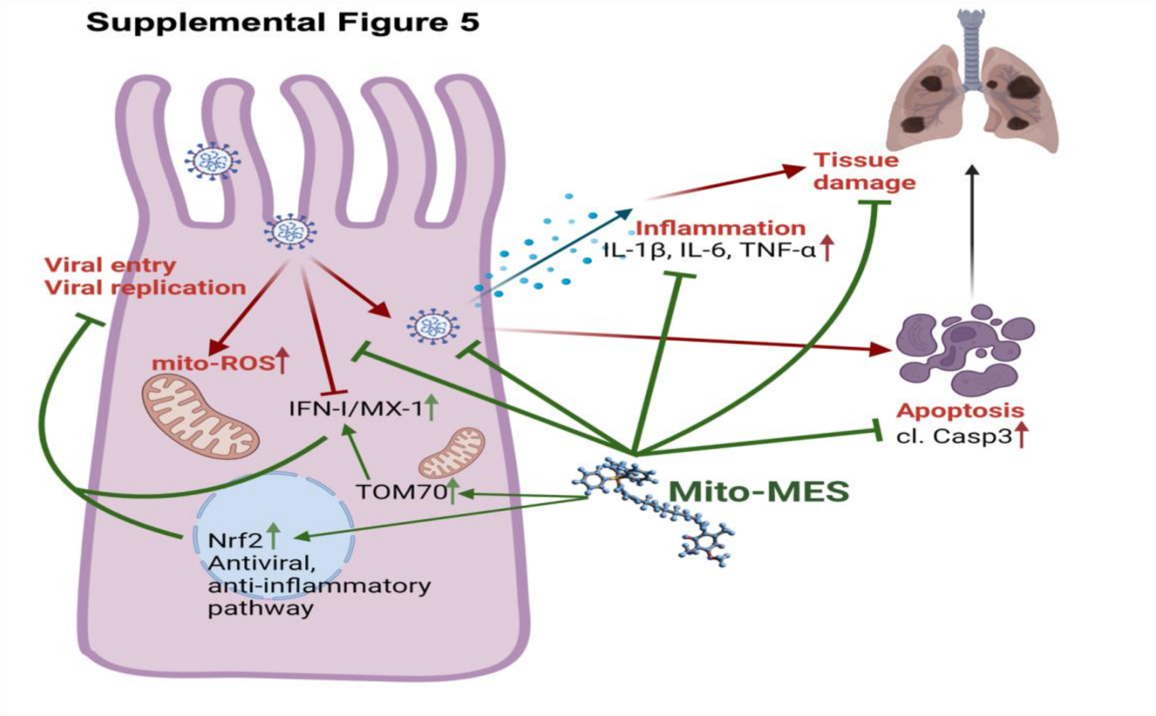


**Fig. S5. Overall hypothesis and impact of Mito-MES on SARS-CoV-2 replication and associated inflammation and tissue damage.** Mito-MES has antiviral, antiapoptotic and anti-inflammatory effect in epithelial cells infected with SARS-CoV-2. Mito-MES has pleiotropic cellular effects that collectively inhibit SARS-CoV-2 replication: 1) blocks SARS-CoV-2 viral entry through its hydrophobic dTPP moiety; 2) inhibits cytoplasmic viral replication through a) its hydrophobic dTPP moiety, b) upregulation of antiviral host pathways (Nrf2 pathway, MX1, TOM70), c) downregulation of mitochondrial reactive oxygen species (mito-ROS) that induce proviral host factors for replication of coronaviruses; 3) blocks SARS-CoV-2-associated cellular apoptosis and subsequent release of SARS-CoV-2 virions to infect other cells. Mito-MES has also anti-inflammatory properties (reduces release of IL-1β, IL-6, TNF-α) that are mediated partially through the Nrf2 pathway. Collectively, the antiviral, antiapoptotic and anti-inflammatory effects of Mito-MES in epithelial cells infected with SARS-CoV-2 reduce tissue damage and morbidity from COVID-19. Created with Biorender.com and Avogadro.cc.
